# KIF5A downregulation in spinal muscular atrophy links axonal regeneration defects with ALS

**DOI:** 10.1101/2025.07.11.664426

**Authors:** Tetsuya Akiyama, Yi Zeng, Caiwei Guo, Olivia Gautier, Lauren Koepke, Juliane Bombosch, Odilia Sianto, Jay P. Ross, Phuong Thi Hoang, Luke Yuchen Zhao, Cole Spencer, Michelle Monje, John W. Day, Aaron D. Gitler

## Abstract

Spinal muscular atrophy (SMA) is a devastating neuromuscular disorder caused by mutations in the *Survival Motor Neuron 1* (*SMN1*) gene, leading to decreased SMN levels and motor neuron dysfunction. SMN-restoring therapies offer clinical benefit, but the downstream molecular consequences of SMN reduction remain incompletely understood. Here, we demonstrate that SMN deficiency results in downregulation of KIF5A in human neurons and in a mouse model of SMA. We provide evidence that reduced SMN levels impair axon regeneration, which is rescued by KIF5A overexpression and that the RNA-binding protein SMN functions to stabilize KIF5A mRNA. These findings provide evidence of a molecular link between SMA and ALS pathophysiology, highlighting KIF5A as a new SMN target. Our findings suggest SMN-independent interventions targeting KIF5A could represent a complementary therapeutic approach for SMA and other motor neuron diseases.

## Introduction

Spinal muscular atrophy (SMA) is the most common genetic cause of infant mortality (1–3). SMA is caused by homozygous loss-of-function mutations or deletions in the *SMN1* gene, which encodes survival motor neuron (SMN) protein. Loss of SMN causes lower motor neuron degeneration, which leads to progressive muscle weakness, paralysis, and eventually death usually within the first two years of life (4). SMN is a highly conserved ubiquitous protein that plays critical roles in multiple cellular processes essential for motor neuron survival and function, including the assembly of small nuclear ribonucleoproteins (snRNPs), pre-mRNA splicing, RNA metabolism, and axonal mRNA transport (5–8).

Humans have a related gene, *SMN2*, which encodes the identical SMN protein and is nearly identical to *SMN1* except for a single nucleotide substitution that affects how the gene is spliced, resulting in exclusion of exon 7 and unstable SMN2 protein (4). Typically, SMA patients have zero functioning copies of SMN1, and the severity of the disease inversely correlates with the copy number of the *SMN2* gene. Humans can have between zero and eight copies of *SMN2* (9), and the more copies of *SMN2*, the milder the clinical features. This observation has led to three different therapeutic strategies, which have significantly improved clinical outcomes by increasing SMN protein levels (10–12). The first approach uses an antisense oligonucleotide (ASO) that blocks the *SMN2* intron 7 splicing repressor, thus improving exon 7 inclusion in *SMN2* transcripts and increased SMN protein levels (13, 14). Another approach delivers adeno-associated viral vectors intravenously to transduce CNS neurons and deliver an episomal *SMN* genetic construct (11, 15) that expresses high levels of SMN protein. Finally, a small molecule delivered orally can block the same *SMN2* intron 7 splicing repressor as the ASO and directly modulate splicing of the *SMN2* gene to boost SMN protein levels (16).

All three of these approaches are now approved by the US FDA and are being used in clinical practice. Simply put, they have changed the natural history for this once fatal disorder (3). But these therapies do not completely eliminate disease-related deficits or prevent their progression, particularly when treatment initiation is delayed or in patients with severe SMA phenotypes (17–19). Moreover, with the splice modulating approaches, there is a natural ceiling effect based *SMN2* pre-mRNA expression levels (20). For gene therapy, there is the potential for long-term toxicity from overexpressing SMN (21), but there are questions of its long term efficacy. Thus, there remains a need to identify complementary therapeutic targets that act independently of SMN restoration. One way to discover such approaches is to define the downstream cellular and molecular consequences of SMN reduction.

Kinesin family member 5A (KIF5A) is a neuron-specific motor protein essential for axonal transport of various cargos, including mitochondria, synaptic vesicles, and mRNAs (22). Mutations in *KIF5A* have been identified as causative factors in adult-onset motor neuron diseases such as amyotrophic lateral sclerosis (ALS) (23, 24), axonal neuropathy (Charcot-Marie-Tooth disease type 2; CMT2) (25), hereditary spastic paraplegia (HSP) (26, 27), and adult-onset distal spinal muscular atrophy (28), highlighting its critical role in maintaining motor neuron integrity. KIF5A-null human motor neurons exhibit impaired axonal regeneration, further underscoring the critical role of KIF5A in axonal repair (29). These findings highlight the importance of axonal regeneration deficits as a common pathogenic mechanism across motor neuron diseases. Despite the established role of KIF5A in motor neuron health, its potential involvement in SMA pathogenesis has not been previously explored.

In this manuscript, while we were analyzing RNA-seq data to investigate the downstream molecular consequences of SMN deficiency, we discovered a dramatic reduction in *KIF5A* expression in SMN-deficient human motor neurons. We confirmed reduced *KIF5A* expression in spinal motor neurons from an SMA mouse model and in SMA patient-derived motor neurons, indicating a conserved pathomechanism. Importantly, we show that restoring KIF5A expression is sufficient to rescue axonal regeneration defects caused by SMN deficiency. We provide evidence that SMN binds and stabilizes *KIF5A* mRNA, suggesting a novel mechanism by which SMN deficiency leads to impaired axonal regeneration. Together, this work reveals KIF5A as a therapeutic target that could complement existing SMN-restoring therapies. Our findings also reveal a molecular link between SMA and ALS, providing new insights into the mechanisms underlying motor neuron degeneration.

## Results

To investigate the molecular mechanisms underlying SMA, we performed RNA-sequencing of human neurons following *SMN* knockdown. We used human neurons derived from either induced pluripotent cells (iPSCs) or human-embryonic stem (ES) cells (Supplementary Table 1) and two distinct systems to induce neuronal differentiation. To make motor neurons, we used HB9-Td-Tomato iPSCs (30), in which Td-Tomato was expressed under the HB9 promoter, allowing for the isolation of motor neurons (iMNs) via FACS (Figure 1A). To generate excitatory glutamatergic neurons, we used H1 ES cells engineered for inducible NGN2 expression (i^3^Neurons; i^3^Ns) (31). We targeted both cell types for *SMN*-knockdown (KD) using lentiviral shRNA against *SMN1/2*. Following 12 days of *SMN*-KD, we harvested RNA for RNA-sequencing (RNA-seq) (Figure 1A).

**Figure 1.**
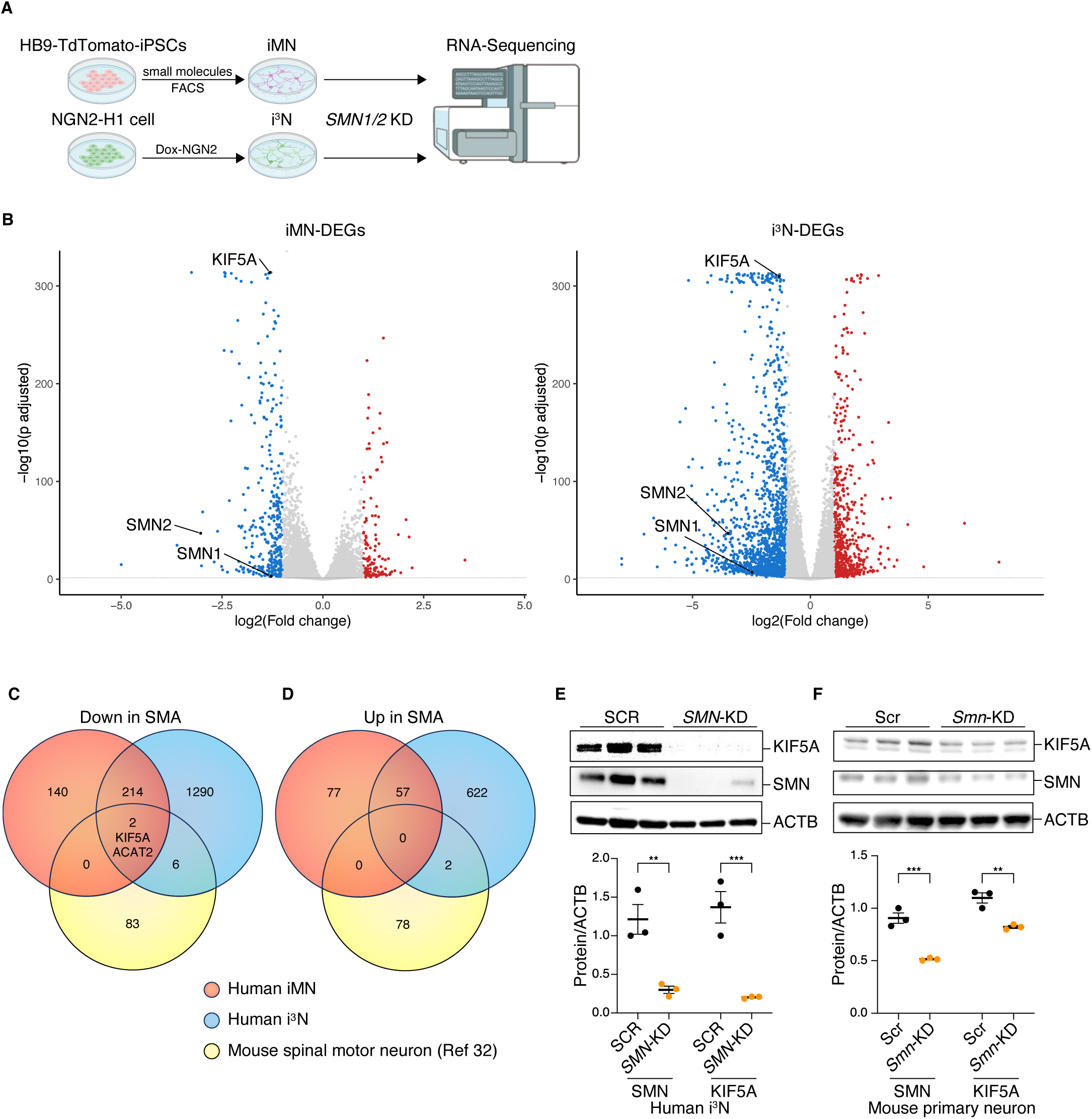
SMN deficiency downregulates KIF5A expression. (A) Two types of human neurons were generated and analyzed. Td-Tomato-positive motor neurons (iMNs) were derived from human iPSCs by small molecule treatment and purified using fluorescence-activated cell sorting (FACS) under the control of the HB9 promoter. In parallel, i^3^Neurons (i^3^Ns) were produced by doxycycline-induced NGN2 expression from isogenic, integrated human H1 ESCs (Supplementary Table 1). Both iMNs and i^3^Ns were subsequently transduced with lentivirus carrying shRNA targeting *SMN1/2* (*SMN*-KD), followed by RNA sequencing. (B) Volcano plots showing differentially expressed genes (DEGs) in iMNs and i^3^Ns following *SMN*-KD. Each plot displays log2 fold change (x-axis) versus negative log10 adjusted p-value (y-axis). Genes significantly upregulated (red dots) or downregulated (blue dots) are defined by adjusted p-value < 0.01 and |log2 fold change| > 1. (C, D) Venn diagrams integrating RNA-seq datasets from iMNs (red), i^3^Ns (blue), and laser-microdissected motor neurons from SMA model mice (yellow) (31). Downregulated DEGs (C) and upregulated DEGs (D) were defined by adjusted p-value < 0.01 and |log2 fold change| > 1. Gene lists are provided in Supplementary Table 2. *KIF5A* was identified as a commonly downregulated gene. (E, F) Western blot analysis of i^3^N lysates following *SMN* knockdown (E), and of primary mouse cortical neurons treated with siRNAs targeting *Smn* (F), confirming the reduction of SMN and KIF5A protein levels. Quantification of SMN and KIF5A protein levels compared to control (scramble; SCR or Scr). Quantification data from three independent experiments are shown for each condition. Statistical analysis was performed using an unpaired two-tailed t-test.

As expected, *SMN1* and *SMN2* RNA levels were significantly reduced (Figure 1B). In iMNs, *SMN*-KD resulted in downregulation of 362 genes and upregulation of 138 genes (Figure 1C, 1D, Supplementary Table 2). In i^3^Ns, 1530 genes were downregulated, and 688 genes were upregulated (Figure 1C, 1D, Supplementary Table 2). Notably, 216 genes were commonly downregulated, and 57 genes were commonly upregulated in both cell types (Figure 1C and D, Supplementary Table 2). To narrow down the list of SMN targets, we compared our set of up-and downregulated genes to a previously reported RNA-seq dataset of motor neurons isolated by laser microdissection from the spinal cord of an SMA mouse model (Figure 1C and D, Supplementary Table 2) (32). There were only two genes, *ACAT2* and *KIF5A*, downregulated in all three datasets (Figure 1C). Immunoblot analysis confirmed that SMN reduction resulted in decreased KIF5A protein levels in human i^3^Ns (Figure 1E), whereas ACAT2 protein levels remained unchanged (Supplementary Figure 1A, B). To confirm whether KIF5A downregulation also occurs in mouse neurons and to assess the conservation of this effect across species, we performed siRNA-mediated *Smn* knockdown in embryonic mouse primary cortical neurons. This resulted in a robust and reproducible reduction in both SMN and KIF5A protein levels (Figure 1F). These findings validate the transcriptomic data and indicate that the downregulation of KIF5A is a conserved and specific response to SMN deficiency.

To validate our in vitro results in vivo, we examined KIF5A expression in the spinal cord of 10-day-old SMNΔ7 mice. These mice carry a homozygous deletion of the endogenous mouse *Smn* gene and are transgenic for one full-length copy of the human *SMN2* gene and one copy of a human SMN cDNA lacking exon 7 (referred to as SMNΔ7). This combination results in a severe SMA phenotype that closely resembles the human disease (33, 34). Compared to age-matched control animals, spinal motor neurons from SMA mice exhibited significantly reduced KIF5A staining (Figure 2). Next, to validate our findings from *SMN*-KD models, we examined KIF5A expression in motor neurons derived from SMA patient iPSCs. We utilized iPSC lines from four healthy donors and six lines from SMA patients (Figure 3A, Supplementary Table 1, which includes patient demographic details, including *SMN2* copy number). We differentiated each cell line to motor neurons. Motor neuron induction efficiency was comparable across cell lines (Supplementary Figure 2). Compared to the four healthy donor lines, all six SMA patient-derived motor neurons exhibited reduced levels of both SMN and KIF5A proteins (Figure 3B). Importantly, upregulation of SMN using a lentivirus was sufficient to restore KIF5A protein levels (Figure 3B), providing evidence that SMN directly regulates KIF5A levels. We also tested an alternative SMN-restoration strategy using the exon 7 splice-skipping ASO, nusinersen, on the SMA patient-derived motor neurons. After 20 days of treatment, we observed a mild but consistent increase in SMN expression and a partial restoration of KIF5A levels (Figure 3C). To further generalize our findings, we reanalyzed publicly available transcriptomic datasets, including a large-scale RNA-seq study of iPSC-derived motor neurons from more than 400 individuals (Supplementary Figure 4A) (35). We found a significant correlation between *SMN* and *KIF5A* expression (Supplementary Figure 4B). This correlation supports the idea that *KIF5A* downregulation is not limited to our specific SMA models. Instead, it likely reflects a broader, disease-relevant mechanism that may impact motor neuron phenotypes across different contexts.

**Figure 2.**
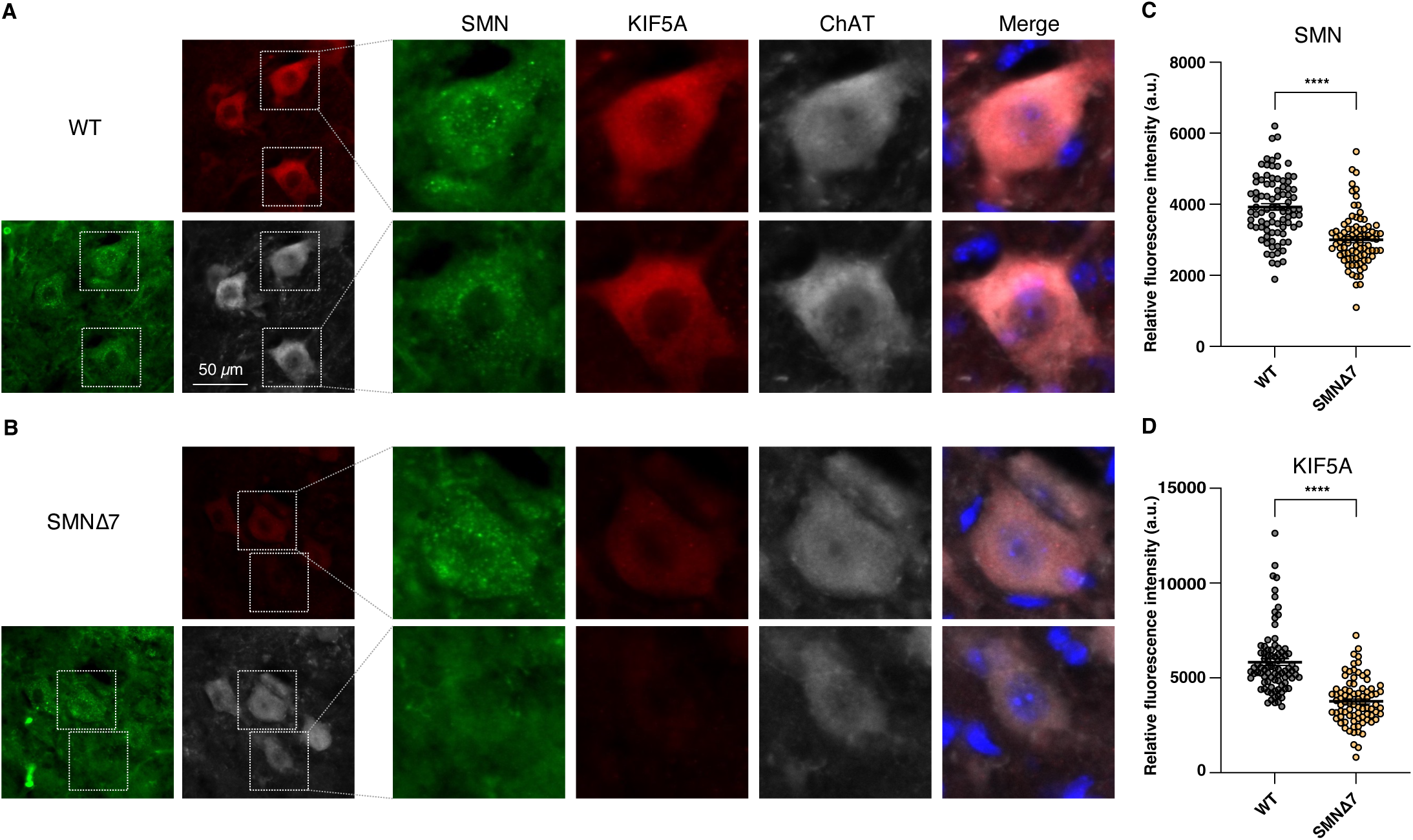
KIF5A is downregulated in SMA model mice. (A) Immunohistochemical analysis of whole spinal cord sections from postnatal day 10 SMNΔ7 mice and age-matched controls was performed. (B) SMA model mice exhibited decreased SMN and KIF5A staining in spinal motor neurons. (C) Quantification for SMN and (D) KIF5A was based on the average of more than 20 motor neurons sampled across non-consecutive sections from 3 biologically independent mice per group. Statistical comparison was performed using unpaired t-test on per-animal averages.

**Figure 3.**
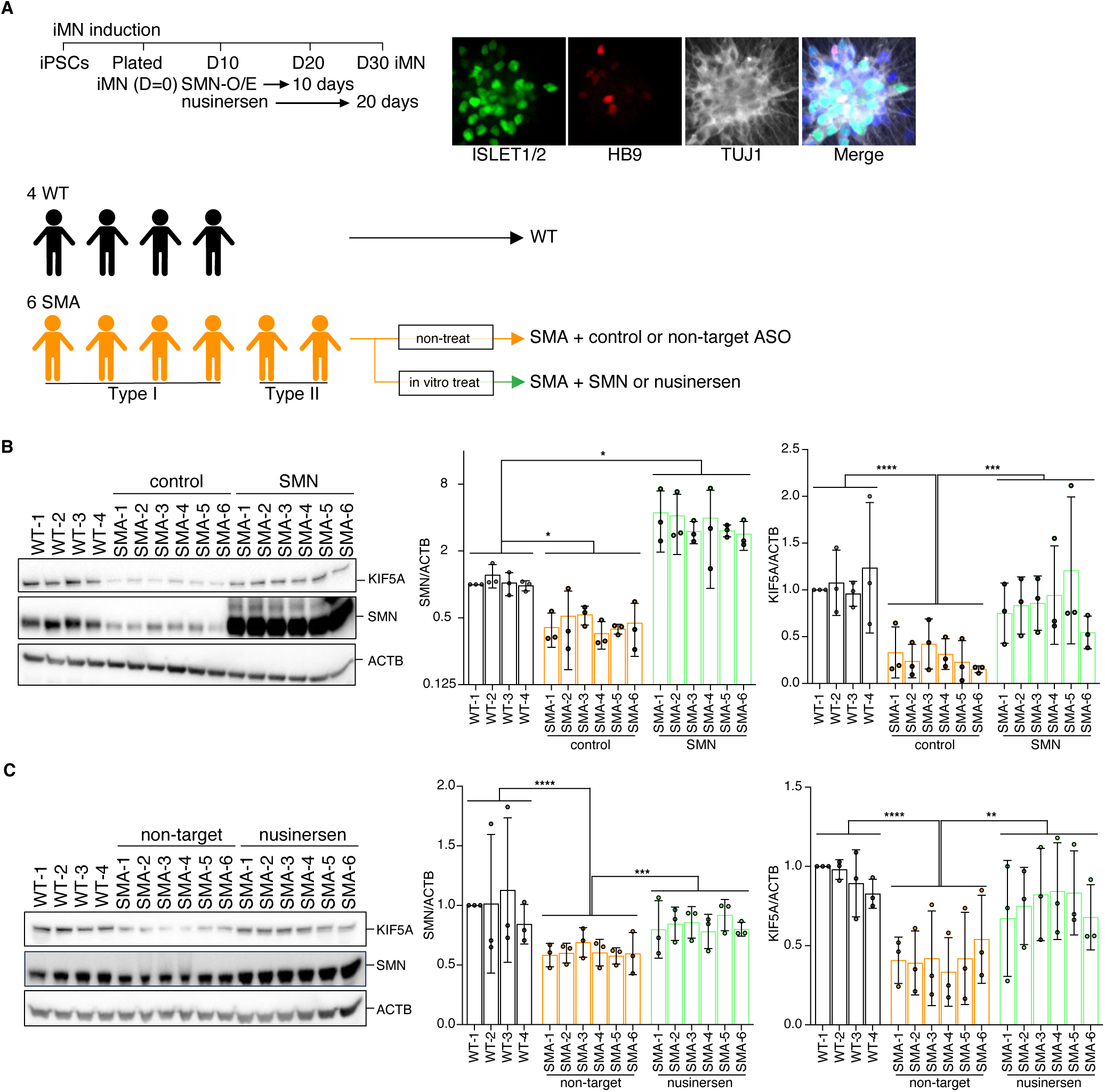
KIF5A is downregulated in SMA patient-derived iPSC motor neurons. (A) Schematic of the experimental design. Motor neurons were differentiated from four healthy donor and six SMA patient iPSC lines using small molecules. SMN overexpression was induced on day 10 after plating, with protein samples collected on day 20. For nusinersen treatment, administration started on day 10 and continued for 20 days; samples were collected on day 30. See Supplementary Table 1 for cell line information. Representative ICC images on day 20 show staining with βIII-tubulin (neuronal marker), HB9, and Islet1/2 (motor neuron markers). Additional data are shown in Supplementary Figure 2. (B) Representative Western blots of SMN and KIF5A in WT and SMA-iMNs transduced with either empty vector (control) or SMN-overexpressing lentivirus (SMN). Bar graphs show SMN/ACTB and KIF5A/ACTB levels normalized to WT1. SMN and KIF5A were reduced in SMA + control, while SMN-OE restored KIF5A to near-WT levels. (C) Western blot analysis of SMN and KIF5A in SMA-iMNs treated with nusinersen or non-targeting ASO. WT samples were included for comparison. Nusinersen increased SMN expression and partially rescued KIF5A levels, though effects were milder and more variable than SMN-OE. All experiments were performed in triplicate and analyzed statistically. Three biologically independent experiments were performed for quantification. Statistical analyses were conducted for each group using one-way ANOVA followed by post hoc Tukey’s multiple comparisons test.

We next investigated the functional impact of KIF5A downregulation in SMA. Given the established role of KIF5A in axonal transport and regeneration (29), we hypothesized that KIF5A downregulation contributes to the axonal growth defects observed in SMA patients and SMA mouse models (36, 37). To test this hypothesis, we performed an axon regeneration assay using *SMN*-KD i^3^Ns. *SMN*-KD significantly impaired axonal regeneration (Figure 4). *KIF5A* deficiency alone also reduced axon regeneration (Supplementary Figure 3A-D), consistent with previous reports (29). Importantly, KIF5A expression provided almost full rescue of the axonal regeneration defect in *SMN*-KD i^3^Ns (Figure 4, Supplementary Figure 3E and F). Thus, KIF5A functions downstream of, or parallel to, SMN in the maintenance of axonal integrity.

**Figure 4.**
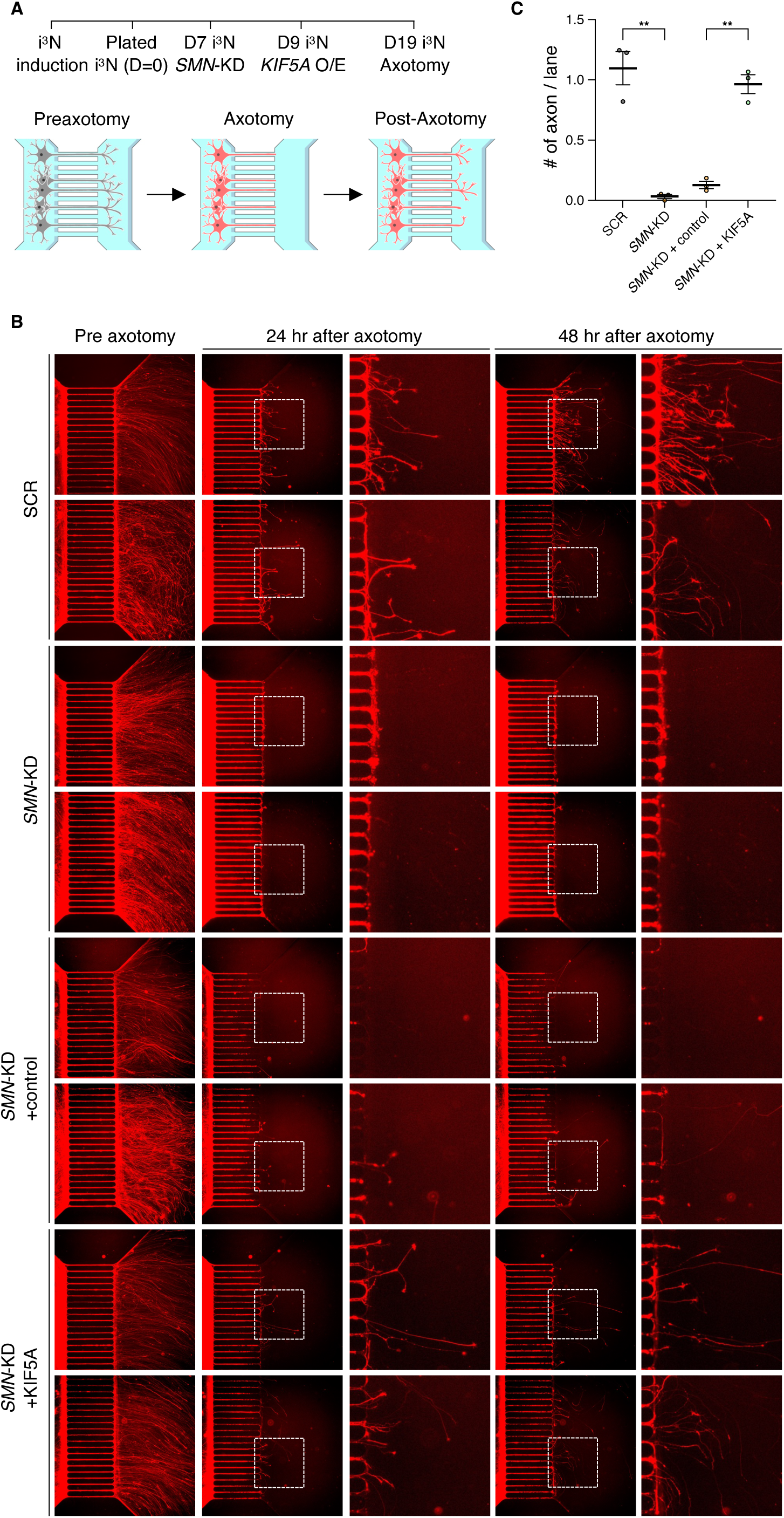
KIF5A overexpression rescues axonal regeneration defects in SMN-deficient neurons. (A) Experimental timeline and schema: i^3^Ns underwent *SMN*-KD for 12 days, followed by axonal transection using a microfluidic device. To overexpress KIF5A in SMN-deficient neurons, cells were infected with lentivirus expressing *KIF5A* two days after initiating *SMN*-KD. Axonal regeneration was assessed by live-cell imaging at 24 and 48 hours post-transection on microfluidic device (XC-450, Xona Microfluidics, see method for detail information). (B) Representative live-cell images before and after axonal transection. Axons extend from the cell body compartment (left) into the axonal compartment (right). The length of each microchannel is 450 μm. (C) Quantification of regenerating axons per microfluidic channel. Data represent the average number of regenerating axons per channel, normalized to control conditions (n = 3 images per condition from two independent experiments; 17–20 channels per image; >100 channels total).

Finally, we investigated the mechanism underlying *KIF5A* mRNA reduction following *SMN*-KD (Figure 5). First, we considered the possibility that SMN deficiency might alter *KIF5A* mRNA splicing. However, re-analysis of our RNA-seq data revealed no clear splicing abnormalities in *KIF5A*. We then hypothesized that SMN deficiency might instead reduce *KIF5A* mRNA stability. To test this, we treated *SMN*-KD i^3^N with Actinomycin D to halt new RNA synthesis and measured *KIF5A* mRNA levels over time by qRT-PCR, using *GAPDH* as a stable control. Compared to control samples (treated with a scrambled, non-targeting shRNA), *SMN*-KD neurons exhibited an accelerated decline in *KIF5A* mRNA levels, indicating reduced mRNA stability (Figure 5B). To determine whether SMN interacts with *KIF5A* mRNA, we performed immunoprecipitation (IP) using an anti-SMN antibody (Figure 2C), followed by RNA extraction (RNA-IP) and PCR analysis. *KIF5A* mRNA was specifically enriched in SMN-IP samples, whereas *KIF5B* mRNA was not detected, suggesting selective binding of SMN to *KIF5A* transcripts (Figure 5D). These experiments provide evidence that SMN associates with *KIF5A* mRNA either directly or indirectly. Together, these results suggest that *KIF5A* mRNA is destabilized in the absence of SMN.

**Figure 5.**
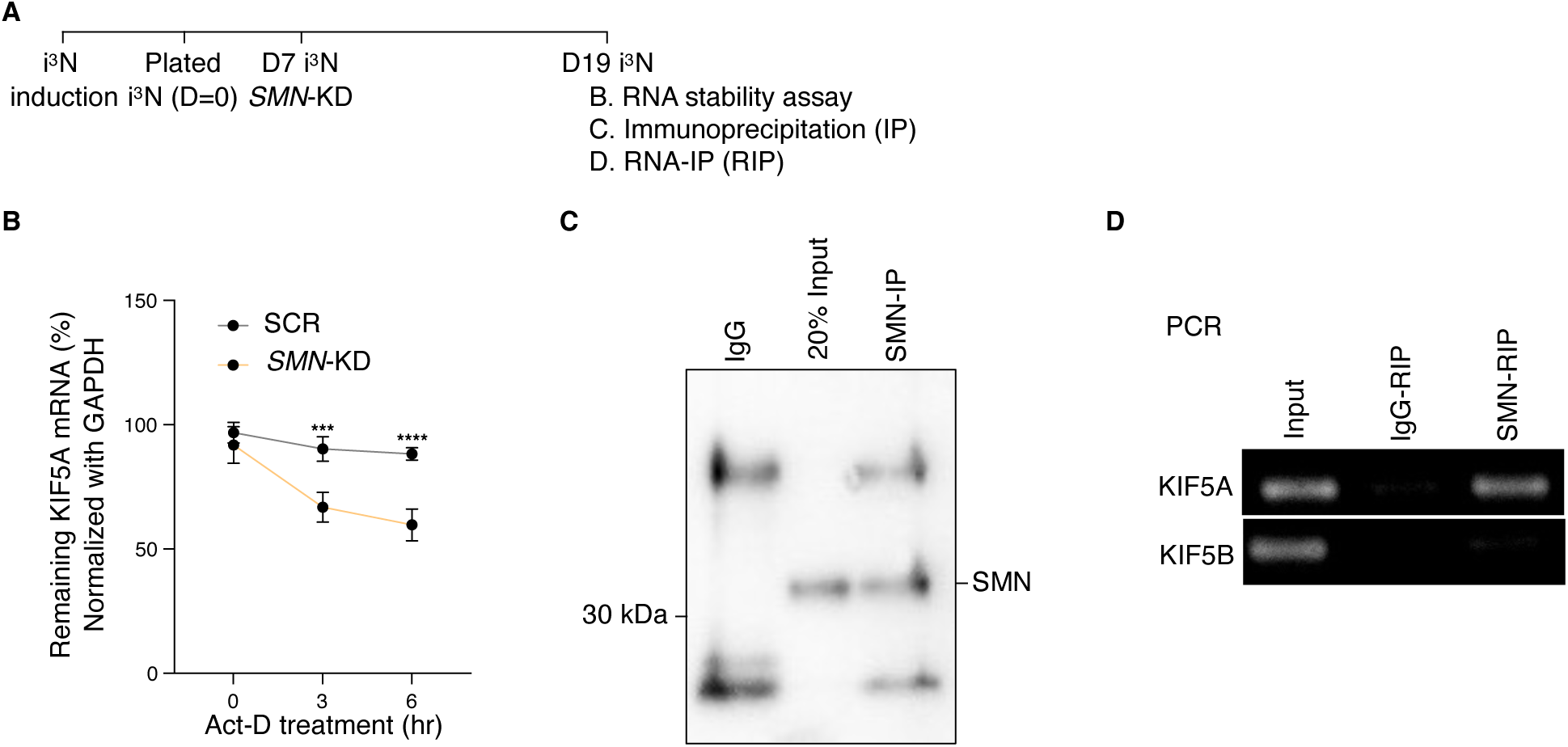
SMN stabilizes *KIF5A* mRNA The relationship between SMN protein levels and KIF5A mRNA stability was investigated. (A) Experimental schematic: 12 days post-induction, *SMN-*KD and SCR i^3^Ns were collected for assays shown in panels B-D. (B) *SMN*-KD and SCR i^3^Ns were treated with Actinomycin D (Act-D, 10 µg/mL) to inhibit new RNA synthesis, and samples were collected at indicated time points for qRT-PCR analysis. KIF5A mRNA levels decreased more rapidly in *SMN*-KD samples compared to SCR controls at 3 hours and 6 hours. Statistical analysis was performed using 2-way ANOVA followed by Šídák’s multiple comparisons test. n = 3 independent experiments. (C) Immunoprecipitation (IP) using an anti-SMN antibody confirmed successful pulldown of SMN protein. (D) RNA immunoprecipitation (RIP) followed by RT-PCR analysis demonstrated enrichment of *KIF5A* mRNA, but not *KIF5B* mRNA, in SMN-IP samples, suggesting specific binding of SMN to KIF5A transcripts.

## Discussion

Recent studies have identified molecular links between SMA and ALS, suggesting shared pathogenic mechanisms involving RNA-binding proteins and spliceosomal dysfunction. For example, interactions between FUS and SMN proteins have been reported, linking ALS and SMA pathogenesis through shared RNA processing pathways (38). Additionally, the ASC-1 complex, implicated in RNA metabolism, has been identified as a molecular link between ALS and SMA (39). Moreover, defects in spliceosome integrity have been observed in both ALS and SMA, highlighting common molecular pathways underlying these motor neuron diseases (40). Our results with KIF5A and SMN now provide an additional link. Furthermore, our analysis of publicly available ALS patient-derived motor neuron RNA-seq data from the Answer ALS dataset indicated a robust correlation between *SMN* and *KIF5A* expression levels, supporting the broader relevance of this pathway across motor neuron diseases (35, 41). Our data extend the known role of KIF5A beyond ALS and HSP (23–28), identifying its dysregulation as a novel contributor to SMA pathogenesis and further supporting a shared mechanism across motor neuron diseases. Thus, KIF5A may serve as a critical molecular link connecting distinct motor neuron disorders through common pathways, underscoring its potential as a shared therapeutic target.

Axonal integrity is crucial for motor neuron function – it facilitates the transport of essential cargoes necessary for neuronal maintenance, growth, and synaptic function. The considerable distance between motor neuron cell bodies and neuromuscular junctions (NMJs) poses unique challenges, making motor neurons particularly vulnerable to disruptions in axonal transport (3, 42). Previous studies have demonstrated that SMN deficiency impairs axonal elongation (36, 37). KIF5A deficiency results in similar axonal regeneration deficits (29). Taken together, these findings suggest that impaired axonal regeneration represents another shared pathological mechanism linking SMA and ALS. Our findings are consistent with these observations and identify KIF5A as a critical downstream mediator of SMN deficiency, whose restoration significantly improves axonal regeneration. Current SMA treatments primarily focus on restoring SMN protein levels, significantly improving clinical outcomes (10, 11, 16). However, despite their clinical success, existing therapies do not fully address all downstream consequences of SMN deficiency, particularly disruptions in RNA processing, axonal transport, and cytoskeletal integrity that contribute to motor neuron dysfunction and degeneration (3, 17, 18). Thus, combinatorial therapies and SMN-independent approaches are actively being explored to enhance therapeutic efficacy (3). Targeting KIF5A could complement existing therapies by directly addressing axonal transport deficits.

Strategies to increase KIF5A levels, such as enhancing mRNA stability, direct protein restoration, or pharmacological activation, could potentially enhance the efficacy of SMN-restoring therapies, particularly in patients with severe SMA phenotypes or those initiating treatment later in disease progression. Future studies should explore the therapeutic potential of these or similar approaches in SMA models to determine their efficacy in restoring axonal transport and improving motor neuron function. Future research should also focus on elucidating the precise molecular mechanisms by which KIF5A influences axonal transport, identifying specific cargoes affected by KIF5A downregulation, and determining the temporal dynamics of these transport deficits in SMA. Additionally, it will be important to investigate whether restoring KIF5A expression or function can ameliorate disease phenotypes *in vivo* using SMA animal models, and to explore the potential for small molecule drugs, antisense oligonucleotides, or gene therapies to increase KIF5A expression or activity in SMA motor neurons. Such studies could provide valuable insights into patient stratification and personalized therapeutic approaches.

In conclusion, our findings identify KIF5A as a previously unrecognized downstream mediator of SMN deficiency, highlighting its critical role in SMA pathogenesis and supporting its potential as a novel therapeutic target. Future studies targeting KIF5A may provide new opportunities for therapeutic intervention, ultimately improving outcomes for SMA patients.

## Materials and Methods

### Stem cell culture and differentiation into motor/i^3^-Neurons

Human iPSCs from both healthy donors and SMA patients (see Supplementary Table 1) were maintained on Matrigel-coated plates in mTeSR plus medium (see Supplementary Table 3 for all medium information) and passaged every 4‒7 days using ReLeSR (30, 43–45). Motor neuron differentiation was initiated using a modified dual-SMAD inhibition protocol (30). Briefly, iPSCs were dissociated with Accutase and seeded as embryoid bodies (EBs) in ultra-low adhesion plates in iMN basal medium (a 1:1 mixture of Advanced DMEM/F12 and Neuro Basal media supplemented with 100 μM ascorbic acid, P/S, Glutamax, NEAA, N2 supplement, and B27 supplement). On Day 0, cells were supplemented with 40 μM SB431542, 0.2 μM LDN193189, and 3 μM CHIR99021. On Day 2, after initial EB formation, the medium was refreshed with the iMN basal medium supplemented with 100 nM retinoic acid and 500 nM smoothened agonist. Subsequent media changes on Days 4, 7, and 9 included additional factors such as BDNF, GDNF, and DAPT (10 μM), and to promote motor neuron specification and maturation. On Day 10, EBs were dissociated using papain-dissociation system and replated onto poly-D-lysine/laminin-coated plates in an iMN basal medium supplemented with BDNF, GDNF, and DAPT. Media were subsequently half-changed every 2–3 days with iMN basal medium supplemented with BDNF and GDNF. For the generation of i^3^N, hES Cell; H1 were used. The differentiation of hESCs into neurons was achieved by inducing the overexpression of NGN2 as previously described (46).

### SMN restoration treatments in iPSC-derived motor neurons

To evaluate the effects of SMN restoration in SMA patient-derived motor neurons, we performed two independent treatment strategies: lentiviral SMN overexpression and nusinersen administration. For lentiviral SMN overexpression, we obtained pLV[Exp]-EGFP-EF1A>hSMN1[NM_000344.4] (Vector ID: VB900104-4718uje) from VectorBuilder. As an empty control, an EGFP-only lentivirus was used. On day 10 after plating, lentivirus was added at a multiplicity of infection (MOI) of 2, and cells were cultured for an additional 10 days before sample collection. For nusinersen treatment, we used a chemically synthesized ASO obtained from MedChem Express (Cat. No.: HY-112980, CAS No.: 1258984-36-9). A non-targeting ASO control was obtained from Qiagen (Antisense LNA GapmeR Standard, 339511). On day 10 after plating, nusinersen and non-targeting ASO was added to the culture medium at a final concentration of 2 µM. The medium was left unchanged for 2 days, after which half-medium changes were performed every 2-3 days for 20 days before sample collection.

### Fluorescence-activated cell sorting (FACS)

Cells were dissociated into single-cell suspensions using a Papain dissociation system and filtered to remove aggregates. Cells were resuspended in sorting buffer consisting of Neurobasal medium (phenol free) supplemented with N2, B27, and ROCK-inhibitor. Sorting was performed using a BD FACS Aria III cell sorter (BD Biosciences) equipped with a 100 µm nozzle. Singlet cells were gated based on forward scatter (FSC) and side scatter (SSC) parameters (FSC-A vs. FSC-H, SSC-A vs. SSC-H) to exclude doublets and aggregates. Cells positive for Td-Tomato were sorted into collection tubes containing culture medium with ROCK-inhibitor and kept on ice until plating.

### shRNA cloning, lentiviral packaging, and cellular transduction

shRNA cloning, lentiviral packaging, and cellular transduction shRNA sequences targeting *SMN1/2* (ATCTGTGAAGTAGCTAATAAT) and scramble control (GATATCGCTTCTACTAGTAAG) were obtained from the Broad Institute Genetic Perturbation Platform (GPP) Portal. Complementary oligonucleotides containing 4-nucleotide overhangs were synthesized, annealed, and ligated into the pRSITER-U6Tet-(sh)-EF1-TetRep-2A-TagRFP vector (Cellecta). Ligated plasmids were transformed into Stbl3 competent cells (Thermo Scientific, C737303) and cultured at 30°C. Large-scale plasmid DNA preparation was performed using the Maxiprep kit (Qiagen), and purified plasmids were used for lentiviral packaging. Second-generation packaging plasmids psPAX2 and pMD2.G (Cellecta) were co-transfected into Lenti-X 293T cells (Takara) using Lipofectamine 2000 (Invitrogen). Lentiviral packaging was conducted at the Stanford Gene Vector and Virus Core (GVVC). Viral supernatants were collected at 48-and 72-hours post-transfection and concentrated by ultracentrifugation. Viral titers were determined by serial dilution and fluorescence-based quantification of RFP-positive cells. Cells were infected with lentivirus at a multiplicity of infection (MOI) of 1. To induce shRNA expression, doxycycline was added to the culture medium. While the standard concentration is 2 μg/ml, a reduced concentration of 0.5 μg/ml was used in this experiment to achieve mild knockdown (Supplementary Figure 1). For *KIF5A*-KD and overexpression experiments, lentiviral vectors were obtained from VectorBuilder. The shRNA lentivirus targeting human *KIF5A* (sh*KIF5A*) was generated using the pLV[shRNA]-mCherry-U6>h*KIF5A*[shRNA#1] vector (Vector ID: VB231128-1302rda), with the target sequence GCGTTGTGAGCTTCCTAAATT. The scramble shRNA control virus was produced using the pLV[shRNA]-mCherry-U6>Scramble_shRNA#1 vector (Vector ID: VB010000-0002vzc). All shRNA lentiviruses were formulated in HBSS buffer and stored at –80°C. For KIF5A overexpression, recombinant lentivirus was produced using the pLV[Exp]-EF1A>hKIF5A[NM_004984.4] vector (Vector ID: VB250107-1212agt). Control virus was generated using the pLV[Exp]-EGFP/Puro-EF1A>mCherry vector (Vector ID: VB010000-9492agg). Both overexpression lentiviruses were formulated in HBSS buffer and stored at – 80°C. Cells were transduced with lentivirus under optimized conditions, and subsequent analyses were performed following transduction.

### Immunoblotting

Cells were lysed in ice-cold RIPA buffer (Sigma-Aldrich R0278) supplemented with a cocktail of protease and phosphatase inhibitors (Cell Signaling Technology, 5872S) at 4 °C for 10 minutes. The lysates were then centrifuged at 20,000 xg for 10 minutes at 4 °C to remove cellular debris, and the resulting supernatant was utilized for BCA protein assay (Invitrogen, 23225) to quantify protein concentrations. Unless otherwise specified, 5-10 µg of protein from each sample was denatured in LDS sample buffer (Thermo Scientific, B0007) containing 10% 2-mercaptoethanol (Sigma-Aldrich) at 80 °C for 10 minutes. The denatured samples were subjected to electrophoresis on 4–12% Bis-Tris Plus Gels (Thermo Fisher, NW04127BOX) and subsequently transferred onto 0.45-μm nitrocellulose membranes (Bio-Rad, 162-0115) using a semi-dry transfer method (Trans-Blot Turbo Transfer System). The membranes were blocked using EveryBlot Blocking Buffer (Bio-Rad, 12010020) or 5% non-fat dry milk in TBS-T for one hour, followed by overnight incubation at 4 °C with specific antibodies against the target proteins (refer to Supplementary Table 5 for details). After triple washing, the membranes were incubated with horseradish peroxidase (HRP)-conjugated secondary antibodies (anti-mouse IgG or anti-rabbit IgG, both at 1:5,000 dilution) for one hour. Detection of protein bands was performed using the Amersham ECL Prime kit (Cytiva, RPN2232), and images were captured using the ChemiDoc XRS+ System (Bio-Rad). Band intensities were quantified using Fiji.

### Immunofluorescence

Cells were fixed with 4% formaldehyde (Electron Microscopy Sciences 15710) in PBS (-) at room temperature for 15 minutes, followed by four washes with PBS (-). They were then permeabilized with 0.2% Triton X-100 in blocking buffer (5% normal goat serum in PBS (-)) at room temperature for one hour. After permeabilization and blocking, cells were incubated overnight at 4 °C in blocking buffer containing specific antibodies (Supplementary Table 5).

Following primary antibody staining, cells were washed four times with 1x PBS and then incubated with secondary antibodies in blocking buffer containing Hoechst 33342 (1:1000) at room temperature for one hour. After this incubation, cells were washed four times with PBS (-) and imaged using a Leica DMI6000 B microscope equipped with a 20x objective and a Hamamatsu ORCA-flash 4.0 camera.

### Total RNA extraction

Total RNA was isolated using RNeasy mini kit (Qiagen) combined with DNase I treatment according to the manufacturer’s protocol.

### qRT-PCR

Total RNA (200 ng) was reverse transcribed into cDNA using the High Capacity cDNA Reverse Transcription Kit (Thermo Fisher, 43-688-13). Quantitative PCR (qPCR) was performed using the PowerUp™ SYBR™ Green Master Mix kit and detected with the QuantStudio 3 Real-Time PCR System (Thermo Fisher). The primers used are listed in Supplementary Table 6.

### Total RNA sequencing

Total RNA from cells treated with scrambled shRNA or *SMN1/2*-targeting shRNA was used to construct RNA-sequencing libraries using a ribosomal RNA depletion protocol (RiboErase), performed by the Stanford Genomics Core Facility. The resulting libraries were quantified, pooled, and sequenced on an Illumina NovaSeq 6000 using an S2 flow cell with 150 bp paired-end reads.

### Gene expression and splicing analysis

Adapter sequences in FASTQ files were trimmed using fastp. The trimmed FASTQ files were then used for transcript quantification with Salmon, followed by differential gene expression analysis using DESeq2, as previously described (46).

For splicing analysis, the adapter-trimmed FASTQ files were aligned to the human genome (hg38) according to ENCODE’s recommended settings using STAR. The uniquely mapped, properly paired reads were subsequently analyzed for splicing events using LeafCutter, which identified cryptic splicing occurrences (47).

### Availability of Data

The sequencing data generated in this study will be deposited in the GEO database.

### RNA Immunoprecipitation (RIP)

RNA immunoprecipitation (RIP) was performed using an anti-SMN antibody and Protein G Dynabeads (Thermo Fisher Scientific, 10004D). RNA was extracted from the immunoprecipitates using TRIzol reagent (Thermo Fisher Scientific, 15596018), and 200 ng of RNA was reverse transcribed into cDNA using the High-Capacity cDNA Reverse Transcription Kit (Thermo Fisher Scientific, 4368813). cDNA was analyzed by RT-PCR using primers specific for *GAPDH*, *KIF5A*, and *KIF5B* (see Supplementary Table 5). PCR products were separated on a 2% agarose gel at 140 V for 20 minutes and visualized using the ChemiDoc XRS+ System (Bio-Rad).

### RNA stability (Actinomycin D) Assay

i^3^N cells treated with sh-SCR or sh-*SMN1/2* were exposed to actinomycin D (10 μg/mL) to inhibit RNA transcription. RNA was extracted at various time points (0, 3, and 6 hours) and analyzed by PCR to assess the stability of *KIF5A* mRNA.

### Axon Regeneration Assay

i^3^N cells were plated at 5 × 10^4^ cells per PDL/Laminin-coated microfluidic device (XC450, Xona Microfluidics). Axons were transected by washing twice with RIPA buffer (10 sec each), followed by three washes each with PBS (-) and i^3^N medium containing 3.3 μg/mL iMatrix-511 (PeproTech, RL511S). Axonal regeneration was quantified by measuring the number and length of regenerating axons at 24 and 48 hours post-transection. Images were acquired using a Leica DMI6000 B microscope equipped with a 10 × objective and a Hamamatsu ORCA-Flash 4.0 camera. For measuring axon length during recovery, axons were counted from a point 5 μm away from the injury site (at the end of the microchannels).

### Image analysis using high content imager

For differentiation efficiency confirmation (Supplementary Fig. 4), ICC-processed cells were obtained from 5 fields/well for D30 neurons using ImageXpress Micro Confocal (Molecular devices), Profiling Ver 4. Imaging conditions were as follows: the nuclei, Islet1/2, HB9, and βIII-tubulin labeled using Hoechst, Alexa Fluor 488, Alexa Fluor 555, and Alexa Fluor 647 antibodies, respectively. Imaging was performed using the following filter set (excitation/emission): nuclei, broad blues (365/535); Islet1/2, greens (475/535); HB9, red (555/595), βIII-tubulin, far reds (630/695). For counting βIII-tubulin positive cell, analysis (Neuronal Profiling Ver 4) began by identifying intact nuclei stained by Hoechst, which were defined as traced nuclei with typical intensity levels lower than the threshold brightness of pyknotic cells. Thereafter, each traced nucleus region was expanded by 50% and cross-referenced with βIII-tubulin. The number of nuclei was analyzed with or without Islet1/2 and/or HB9 staining at the nucleus of βIII-tubulin positive cells.

### Primary mouse cortical neurons

Mice were bred and used as approved by the Administrative Panel of Laboratory Animal Care (APLAC) of Stanford University, an institution accredited by the Association for the Assessment and Accreditation of Laboratory Animal Care (AAALAC). Primary mouse cortical neurons were dissociated into single-cell suspensions from E16.5 mouse cortices using a papain dissociation system, as previously described (48). Neurons were seeded onto PDL-coated plates and grown in Neurobasal medium supplemented with B-27, GlutaMAX, and penicillin– streptomycin (See Supplementary Table 4 for detailed medium information). Half-media changes were performed every 4–5 days. Neurons were plated in 24-well plates (350,000 cells/well). Three days after plating, primary neurons were treated for 3 days with 100 nM-siRNA targeting *Smn* (Dharmacon, L-044280-00) or a non-targeting control (Dharmacon, D-001810-10) using DharmaFECT 3 (Dharmacon, T-2003) in Opti-MEM buffer.

### Mouse immunohistochemical analysis

SMNΔ7 model mice (Jackson Laboratory) were used. Ten days after birth, age-matched mice were sacrificed, and fresh spinal cords were extracted as following. Anesthetized mice were perfused with PBS and the spinal cords were carefully dissected and washed in chilled PBS. The spinal cords were fixed with 4% PFA in PBS at 4°C for 48h and then stored in 30% sucrose in PBS. Fixed spinal cords were mounted in OCT and dissected into 50 µm thick sections using a Leica CM3050 S Cryostat with a cryo-microtome (Leica) and mounted on slides, which were stored at -80°C until used for immunohistochemistry (IHC). For IHC, each sample was washed twice with PBS (-), and blocked with 5% NGS, 1% BSA, and 0.4% Triton-X in PBS (-) for 1 hour at room temperature. The first antibody (see Supplementary Table 5) was applied and incubated at 4°C overnight. After incubation with the first antibody, samples were washed three times with PBS (-) and stained with the second antibody for 1 hour. They were then washed three times with PBS (-), mounted with ProLong Gold (Thermo Fisher Scientific, P36931) overnight, and captured using a microscope. All images were taken under the same conditions and analyzed with Fiji. For analysis, motor neurons were identified using ChAT staining. By applying a threshold, the ChAT-positive area was recognized as the motor neuron area (region of interest, ROI). After analyzing ChAT intensity, the same ROI was applied for SMN and KIF5A staining, and each intensity was calculated (see Supplementary Figure 5).

### Statistical Analysis

Statistical analyses were performed using GraphPad Prism (version 10.4.1). For comparisons, one-way ANOVA followed by Tukey’s post hoc test was used unless otherwise noted. Statistical significance was defined as follows: *p < 0.05, **p < 0.01, ***p < 0.001, ****p < 0.0001.

## Ethics Statement

All experiments involving human iPSCs and ESCs were conducted in accordance with Stanford University’s Stem Cell Research Oversight (SCRO) guidelines and approved under protocol SCRO-858. All recombinant DNA work, including vector cloning, lentiviral packaging, and bacterial manipulations, was approved by the Stanford Administrative Panel on Biosafety (APB) under protocol APB-5676. Mouse experiments are approved by the APLAC of Stanford University, an institution accredited by the AAALAC.

## Acknowledgements

We thank members of the Gitler lab for helpful discussions and comments on the manuscript. We also thank the Wichterle lab for sharing the HB9 reporter line. We thank the NIGMS Human Genetic Cell Repository at the Coriell Institute for providing the WTC-11 iPSC line (GM25256), originally submitted by Dr. Bruce R. Conklin (Gladstone Institute of Cardiovascular Disease, UCSF). The human iPSC lines, GSB-L480 and GSB-L2394, were obtained from Greenstone Biosciences, Inc. T.A. is supported by NIH (2T32AG047126-06A1) and a fellowship from the Takeda Science Foundation. Y.Z. is supported by a postdoctoral scholar award from The Phil and Penny Knight Initiative for Brain Resilience at the Wu Tsai Neurosciences Institute, Stanford University, and a fellowship grant from the Larry L. Hillblom foundation. C.G. is supported by Milton Safenowitz Postdoctoral Fellowship Program. J.P.R is supported by an ALS Scholars in Therapeutics award from the Sean M. Healey & AMG Center for ALS, ALS Finding a Cure, and Fight MND. J.W.D is supported by SMA Foundation. M.M. is supported by SMA Foundation. A.D.G. is supported by NIH (grants R35NS137159, U54NS123743, R01AG064690), Target ALS, and SMA Foundation. A.D.G. is a Chan Zuckerberg Biohub – San Francisco Investigator. To evaluate iMN induction efficiency, all images and analysis were conducted with Stanford High-Throughput Screening Facility (HTS). Lentiviruses were produced at the Stanford Gene Vector and Virus Core (GVVC).

## Author contributions

T.A. and A.D.G. conceived and supervised the study and wrote the manuscript. T.A. designed and performed the experiments and analyzed data. Y.Z. and C.G. performed RNA-seq data analysis. O.G. and L.K. prepared mouse tissue samples for analysis. J.B. prepared primary mouse neurons. O.S. generated and maintained i^3^Ns. J.P.R. analyzed public RNA-seq datasets.

P.T.H. established the iMN induction system. L.Y.Z. and C.S. assisted with ICC and western blot experiments. M.M.D. and J.W.D. supervised parts of the project, contributed to experimental design and interpretation.

---

**Supplementary Figure 1.**
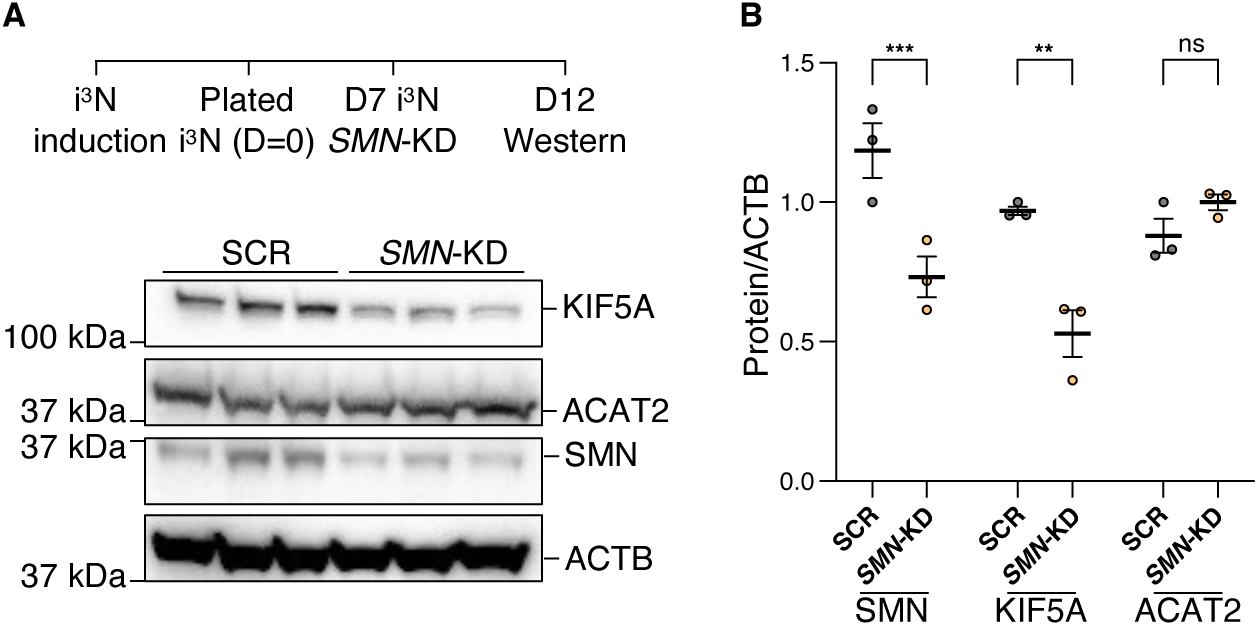
SMN knockdown decreases KIF5A but not ACAT2 levels in i^3^Ns. (A) Knockdown of *SMN* in i^3^Ns led to reduced levels of both SMN and KIF5A proteins, whereas ACAT2 levels remained unchanged. To achieve a mild knockdown, the concentration of doxycycline was reduced to 0.5 μg/ml (standard concentration = 2 μg/ml). (B) Quantification of SMN, KIF5A, and ACAT2 protein levels compared to control shRNA (scramble; SCR). Statistical significance was determined using Student’s t-test. SCR: scrambled shRNA-treated samples; *SMN*-KD: *SMN* shRNA-treated samples. n = 3 independent experiments.

**Supplementary Figure 2.**
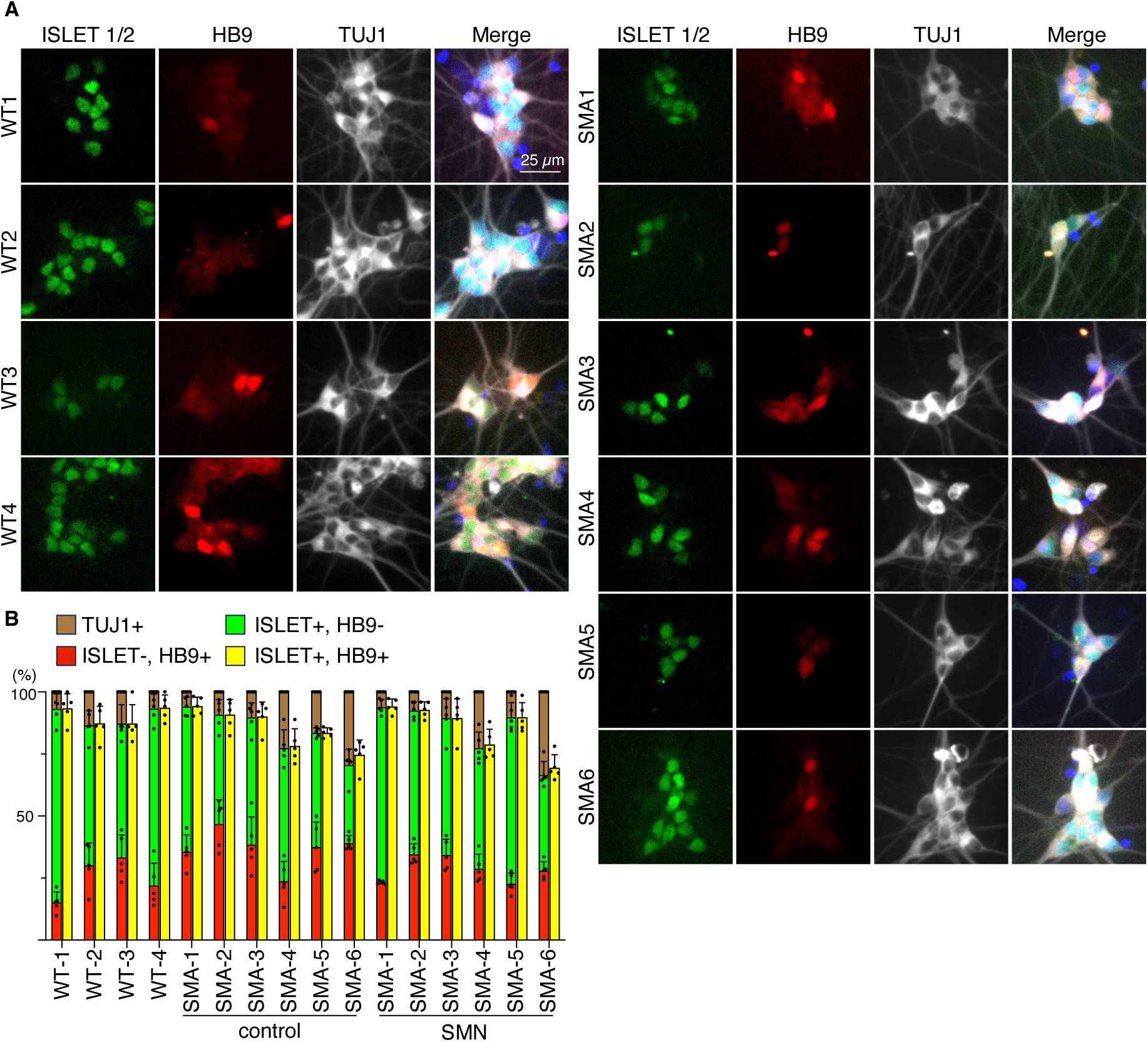
Motor neuron differentiation efficiency is consistent across all iPSC lines. To confirm comparable motor neuron differentiation efficiency between wild-type and SMA lines, immunostaining was conducted on day 20. (A) Representative immunofluorescence images of iPSC-derived motor neurons at day 20, stained with βIII-tubulin, HB9 (red), and ISL1/2 (green), both of which are motor neuron-specific markers. (B) Quantification of induction efficiency was performed using the ImageXpress system. Five regions per sample were imaged, and five independent samples were analyzed to generate the bar plot. In the bar graph, brown indicates the total population of TUJ1-positive nuclei (100% reference). Red and green represent nuclei positive for HB9 or ISL1/2 alone, respectively. Yellow indicates the proportion of nuclei positive for either HB9 or ISL1/2. Statistical analysis was performed using One-way ANOVA on the aggregated values for TUJ1 positive, HB9 positive, ISL1/2 positive, and HB9+ISL1/2 positive nuclei across groups, and no significant differences were observed.

**Supplementary Figure 3.**
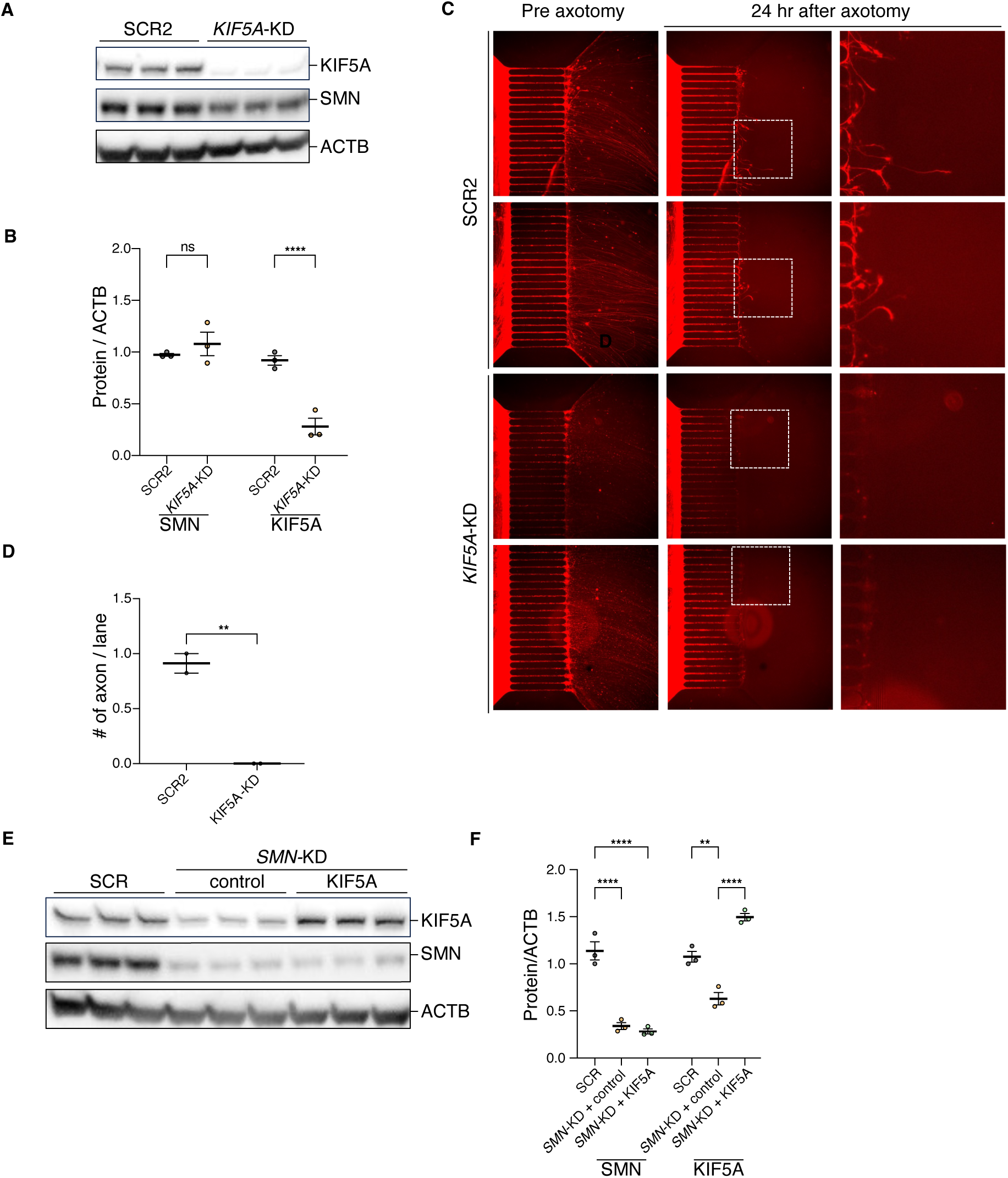
*KIF5A*-KD impairs axonal regeneration. (A, B) Western blot analysis confirming *KIF5A*-KD i^3^Ns after 10 days. The p value was calculated by Student’s t test; SCR, samples treated with scrambled shRNA; *KIF5A*-KD, samples treated with *KIF5A*-shRNA. (C) Representative live-cell images before and after axonal transection. Axons extend from the cell body compartment (left) into the axonal compartment (right). The length of each microchannel is 450 μm. (D) Quantification of the number of regenerating axons per microfluidic channel. Data represent the average number of regenerating axons per microfluidic channel, normalized to control conditions (n = 3 images per condition from two independent experiments, 17-20 channels per image, >100 channels total). Statistical analysis was performed using two-way ANOVA. (E, F) Western blot analysis confirming KIF5A overexpression in *SMN*-i^3^Ns. The p value was calculated by 2-way ANOVA, followed by post-hoc tests.

**Supplementary Figure 4.**
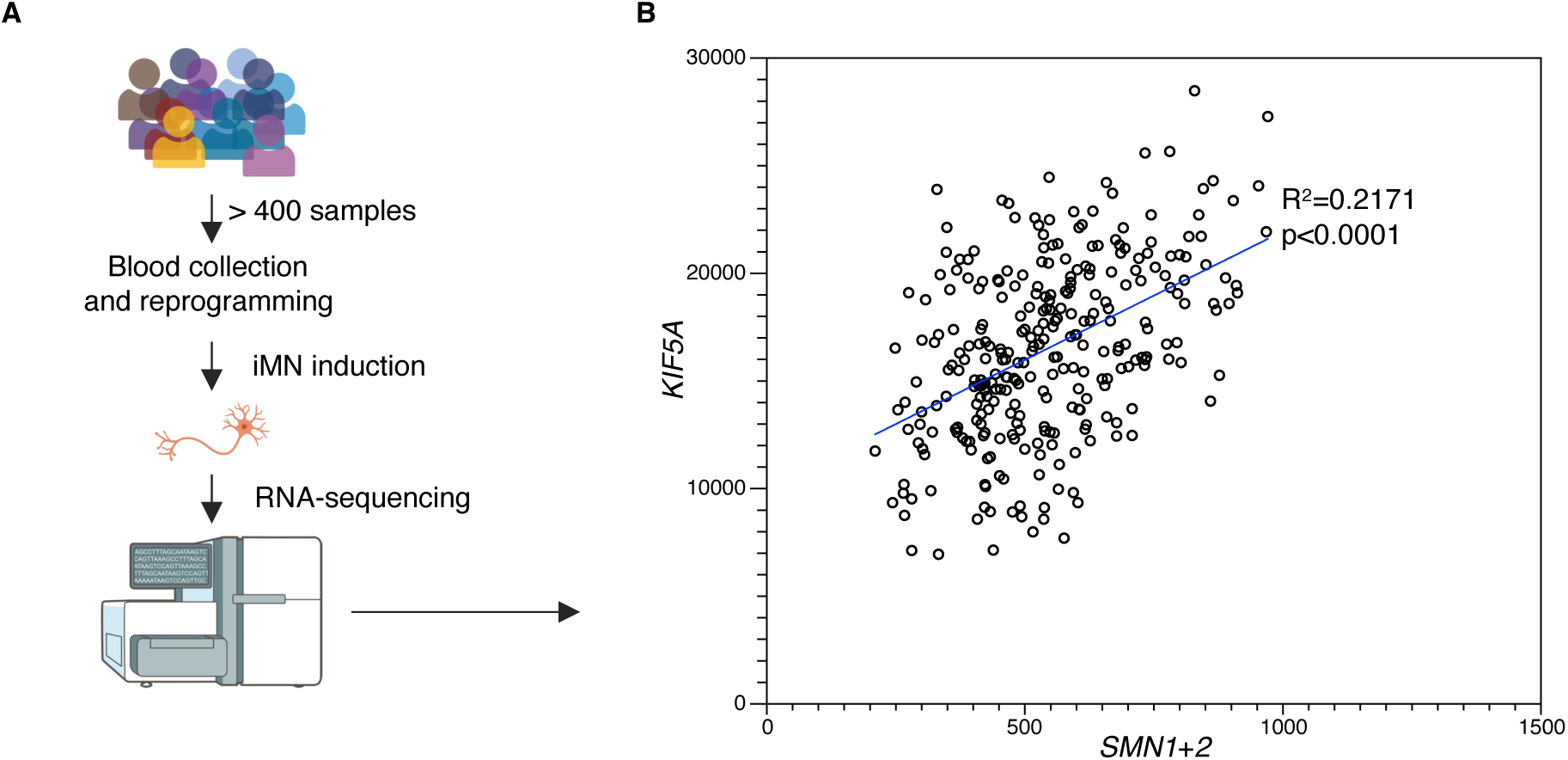
Correlation of *KIF5A* and *SMN1+2* expression in answer ALS iMN dataset. (A) Schematic depicting how RNA-seq data were generated. (B) Scatter plot showing the correlation between *KIF5A* and (*SMN1*+ *SMN2*) expression in iMNs from the Answer ALS dataset. Pearson correlation coefficient and p-value are indicated.

**Supplementary Figure 5.**
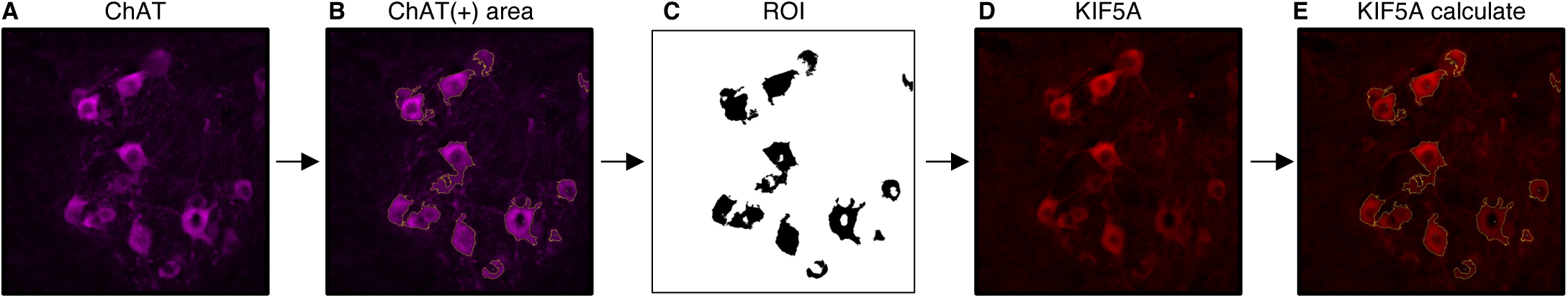
Quantification strategy for spinal cord immunohistochemistry (IHC) Motor neurons were identified using ChAT (Far red) staining (A). By applying a threshold (B), the ChAT-positive area was recognized as the motor neuron area (region of interest, ROI) (C). After analyzing ChAT intensity, the same ROI was applied for SMN (not shown) and KIF5A (D) staining, and each intensity was calculated (E).

